# Redlisting the redlist: *a global analysis of species distributions and biodiversity*

**DOI:** 10.1101/676460

**Authors:** Alice C. Hughes

**Affiliations:** Centre for Integrative conservation, Xishuangbanna Tropical Botanic Garden, Chinese Academy of Sciences

**Keywords:** IUCN, Amphibians, Mammals, Range, global distributions, biodiversity, conservation, reassessment

## Abstract

Global conservation and research has come to rely on the IUCN species range-maps to direct research, allocate funding and define and design protected areas. However, IUCN species range-maps may be created on the basis of little actual data. Though the IUCN methods may be suitable for well known species, they may be liable unreliable for lesser known species or areas. In such cases, human biases may limit the usefulness of the output maps and potentially misdirect conservation funding and protection. Possible errors in these IUCN maps has global implications for the preservation of biodiversity, as flawed data leads to flawed decisions, which are of critical importance to the protection of the biodiversity of this planet. Ultimately we show that the current IUCN “expert-assessment based” range-maps may hinder rather than assist global conservation, as errors ranged from tens to thousands of kilometres (between recorded distributions and IUCN-maps), and for some taxa under 50% of records fell within their mapped-ranges. IUCN maps are being used globally to evaluate global conservation and protection, yet up to 85.4% of IUCN species-range boundaries follow political-boundaries, making using IUCN maps to evaluate the protected-area system and it’s efficacy impossible. The current availability of data, methods and computer-power has relegated these maps in the wake of more empirically based methods, and better use of available data provides the ability to make better conservation decisions, especially for highly threatened species and regions.

## Introduction

The IUCN species range-maps provide a useful basis for conservation decisions, on regional to global scales, as a reference to researchers at all stages and as a basis for allocating grants and protecting and prioritising regions. Indeed the IUCN website (www.iucnredlist.org..../iucnspatialresources) states the purpose of the dataset is “to inform conservation planning and other decision making purposes’, and these global distribution maps have come to replace some of the former regional databases of faunal distributions (i.e. the Southeast-Asian mammal Databank: http://www.ieaitaly.org/samd/). Furthermore the range-maps also provide a source of data for global analyses of species distributions, gap analysis of protected areas and even mapping biodiversity, which shapes decisions and understanding of the distribution of biodiversity and species across the planet.

However, “with great power, comes great responsibility” and thus, though these maps have the power to facilitate effective conservation measures, any errors have the potential to seriously undermine effective conservation for species globally. All analyses and decisions which use IUCN maps are made on the basis that the IUCN range-maps provide a reliable reference source for species distributions, yet these maps are developed by teams on the basis of “expert knowledge” and thus like many forms of analysis which depends on human knowledge rather than empirical data they are subject to bias.

Errors in these maps has the potential to ignore areas of high conservation value, and lead to their under-protection and increase the hardship in attaining funds whilst simultaneously protecting areas which may have fewer, or less unique species. Furthermore by overestimating species ranges it is impossible to reliably gauge population or vulnerability, and thus the assigned species IUCN-redlist status may be in a lower risk category than actually warranted, meaning lower levels of protection and less access to funding than the species deserves. Ultimately any errors in the IUCN maps has the potential to negatively effect conservation planning and management on a global scale, as flawed data may ultimately lead to flawed decisions.

Conservationists and ecologists may dismiss these potential errors as unimportant, however to do so is to underestimate the use and the implications of the use of these range-maps. Indeed a search for the term “IUCN range-maps” on google-scholar results 40800 results, including a range of journals of all impact-factors. A search for ‘IUCN range-maps + global analysis’ yields 30600 results, indicating the use of these maps in a variety of forms. Some of the more significant uses of these maps include a “Global gap analysis: priority regions for expanding the global protected-area network” (Rodrigues et al. 2004), the “global distribution and conservation of rare and threatened vertebrates” (Grenyer et al. 2006), Global patterns of Terrestrial diversity and conservation (Jenkins et al., 2013) and “The performance of the global protected area system in capturing vertebrate geographic ranges” (Cantú-Salazar et al. 2013), and “Global protected area expansion is compromised by protected land-use and parochialism” (Montesino Pouzols et al. 2014). Thus these maps provide a convenient data source to develop a global protected area network, in addition to prioritising regions and managing areas to promote biodiversity retention, and as a result if these maps are incorrect it will have an effect on all decisions based on that data.

Here we use two decades of globally available data from the global-biodiversity information facility (www.gbif.org), this has been extensively cleaned and validated (giving 668583 unique records) before being used to assess the match between empirical data on 5747 species of mammal and amphibian with their IUCN maps.

To make effective decisions requires accurate information. Thus we evaluate the accuracy of global IUCN maps for 3040 mammal and 2707 amphibian species, and assess their merit relative to other methods to ensure that important decisions are made using the best available information, and provides sounder basis for conserving global biodiversity.

## Methods

IUCN range-maps for all amphibians and mammals were downloaded from the IUCN redlist site (http://www.iucnredlist.org/). All distribution point data for mammals and amphibians was downloaded by family for the last two decades from GBIF (http://www.gbif.org/), in addition we used a previously compiled bat database (Hughes et al., 2011, 2012), and the African bat database (http://www.africanbatconservation.org/) (as accurately identified bat specimens are often rare in museum collections). Data was then extensively cleaned in excel (see Appendix S1), any genus unassigned to species was classified based on the longitude and latitude given relative to others of the genus (within 0.1° for both parameters), and where little probable ambiguity existed. Species forming a number of discrete distributional units and at least five decimal degrees from other populations were classified as sub-groups and examined relative to remaining distribution data and IUCN range polygons separately, to remove captive individuals and alien releases of species from the database.

3040 mammal species (based on 512100 records) and 2707 amphibian species (records) were considered overall. For each mammal taxa: 1108 bat species (140287 points), 927 Rodentia (55081 points) 141 Certartiodactyla, 161 carnivores, 50 Dasyuromorphia, 5 Hyracodica, 11 Perissodactyla, (165316 points) 29 Afrosorcida, 13 Cingulata, 196 Eulipotypha, 8 Pilosa, (23659 points), 53 Lagomorpha, 5 Pholidota, one Tubulidentatia (33600 points), 2 Dermoptera, 188 Primates, 8 Scandentia, (1727 points), and two Proboscidea (252 points) in addition to 179 Marsupials and three Monotremata (125778) were considered in analyses, though other species for which locality data existed had not been mapped by the IUCN. For amphibians, 2314 anuran species (132647 points), 41 Gymnophiona (164 points) and 352 Caudata species (23672 points) were considered, See Appendix S1 for supplementary methods.

### a). Range coverage of IUCN maps

Once distribution data for each taxa had been cleaned (see Appendix S1) ArcView 10.1 was used to spatially overlay the point data with the IUCN range-maps for the appropriate taxonomic group. The tabulated intersection of the data was then exported to excel for further processing to determine the percentage of points for each species fell within the appropriate range-map. We also mapped, for all placental mammals (most marsupial groups are only found within a single landmass) what degree of error existed in various “biogeographic” zones (North America, South America, Europe, Eurasia, Africa, Southeast Asia and Australasia) for each mammalian and amphibian order. Europe and Eurasia were separated on the basis that Europe is largely densely populated and many species are intensively studied, whereas Eurasia is a much large area, with lower population density and less ecological research.

### b). Spatial disparity between recorded distributions and IUCN species range maps

We also calculated the degree of spatial mismatch between actual species distribution records and the IUCN range-maps. The maximum and minimum latitudes and longitudes of for each species both from actual data and from the IUCN range-maps were extracted, and the degree of congruence compared. As this analysis is global and a large number of species are distributed between all four hemispheres it is only possible to ascertain the minimum distance between the limitations of a species range based on point data and those of the IUCN ranges, but not if those limitations lay within or outside the mapped ranges. This is because in Western and Southern hemispheres negative values denote that species actual recorded distributions fell within the species range, whereas in Eastern and Northern hemispheres positive values denote that species recorded localities fell within the mapped range. Because species mapped to cross between hemispheres do not always have the data to support such a trend, and some species recorded in a single hemisphere have recorded localities which do cross between hemispheres it is not possible to really separate. Obviously discrepancies between mapped are recorded localities do not necessarily mean the range-map exceeds the species actual distribution, as recorded distribution points are unlikely to indicate the edge of a species range (especially for species with few records), however using spatial mismatch of boundaries in combination with the proportion of localities fell within mapped ranges gives two measures off differences between actual and mapped distribution.

### c). Species boundaries and political boundaries

The concordance of species range boundaries and political boundaries were examined for the six mammal orders which spanned at least five of the biogeographic zones, giving a total of 4272 mammal species (2119 Rodentia species, 1141 Chiroptera species, 442 Eulipotyphla, 246 Carnivora, 232 Cetartiodactyla and 92 Lagomorphia). To examine the concurrence between political boundaries and species ranges the IUCN range-maps for each species were converted to points using the “vertices to points” tool in ArcGis (producing upto 7803124 points for each of the six orders). Using the country boundary data (http://www.diva-gis.org/Data) the spatial join function of ArcGis was used to intersect each species range-map with any nearby political boundaries. A 25km buffer each side of the political boundary was specified, to capture the ranges of any species which “approximately shadowed” a political boundary. The process was repeated using the output of the political boundary intersection with a coastal boundary (http://www.ngdc.noaa.gov/mgg/shorelines/gshhs.html). The total number of points which formed each species range boundary was compared to the number along the political boundary, the coastal zone and related measures to determine the degree to which the political boundary was used as the range delimiter of any given species. The results were then broken down by order and by region, to look for spatial biases in the usage of boundaries to define range limits. Zones with relatively few political boundaries (i.e. Australasia) are unlikely to utilise these boundaries heavily, whereas areas such as Europe which have a relatively large coastal zone will reflect the usage of coastal to non-coastal political boundaries through the proportionate discrepancy between total non-coastal political boundary and percentage of non-coastal range-boundary which overlaps with a political border.

## Results

### a). Range coverage

For the 3040 mammal species considered, 78.59% of recorded localities overall fell within species range-boundaries. Species with few recorded localities, and small orders often had 100% of localities within the IUCN range for each species considered (*Dermoptera, Paucituberculata, Pholidota, Tubulidentata*). Scandentia was the least well mapped taxa, with only 23.35% of recorded localities falling within IUCN ranges, followed by Certartiodactyla with 52.57%. Chiroptera was the next most poorly mapped taxa (Figure 1-2), with 63.95% of records falling within the appropriate ranges, however it should be noted that over 100 bat species with locality data have not been mapped by the IUCN. Only one placental family with more than five species had over 80% of recorded localities falling within the appropriate range-maps (Pilosa: 85.6%), though marsupial species were considerably better, with two families (both with 12 species) showing over 80% congruence between recorded localities and mapped range (Peramelemorphia 81.78%, Macroscelidea 83.4%) and a number of smaller families also showing over 80% congruence (Microbiotheria: 90.32%, Notoryctemorphia: 96.72%). The Monotremata also showed a high level of congruence with 99.3% of localities falling within species mapped ranges (Table 1). A full breakdown of accuracy by family can be found in Appendix S2 (Commentary).

**Table 1.**
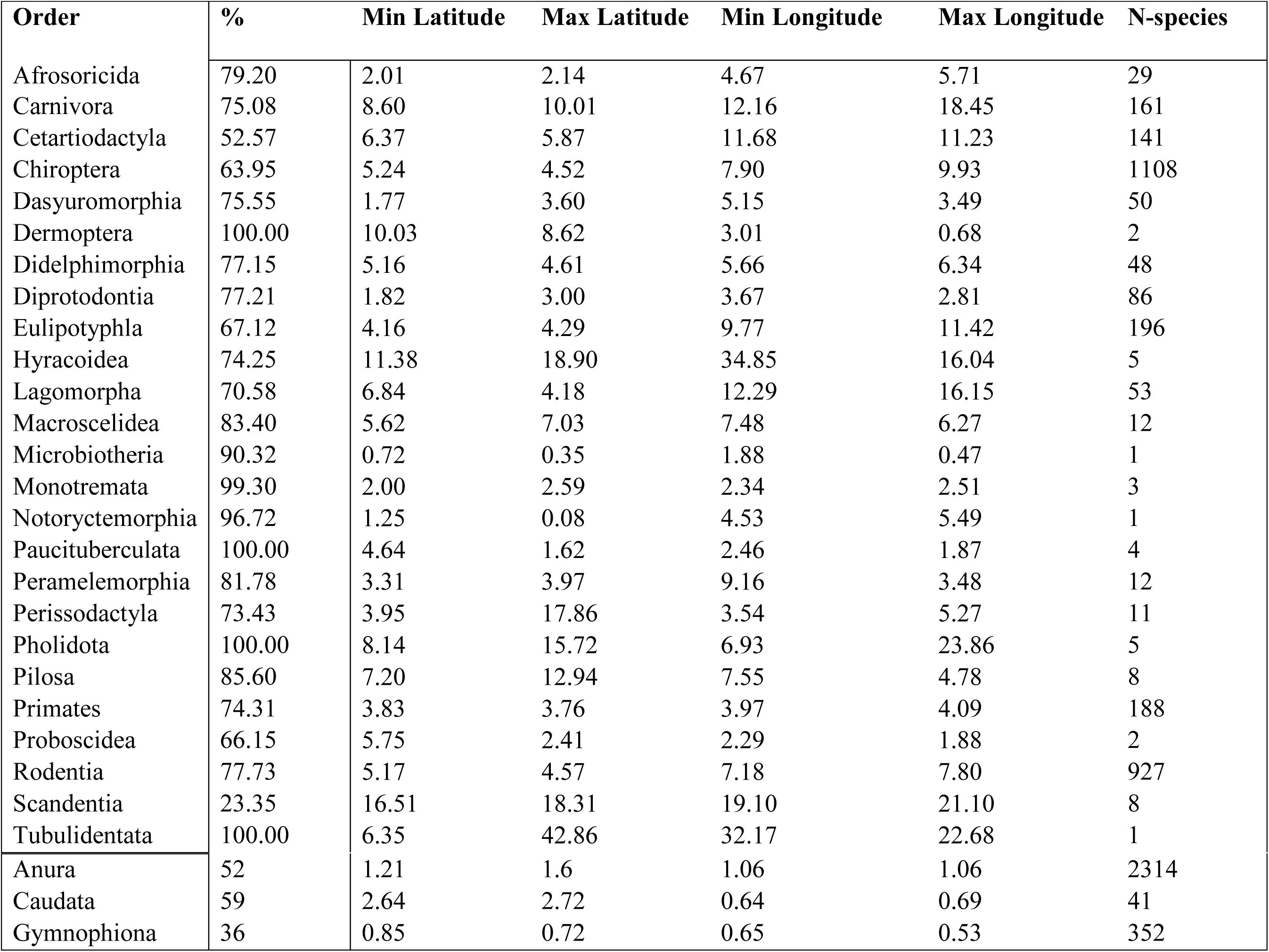
Accuracy by family of the IUCN range maps, mammals are shown at family level and amphibians at order (data for each family is available in Table S1). The first column shows the percentage of recorded localities fall within the appropriate range map for each species within that family. The following columns show the mean distance between the recorded range boundaries and those showed by IUCN range maps. A this analysis combined data from all four hemispheres all minuses were removed (i.e. the minus sign itself was removed after the discrepancy distance between recorded location boundary and mapped boundary had been calculated) before the average for each family was calculated. The number of species within each order considered in this analysis is also listed. Orders aligned to the left are mammals, whereas those on the right are Amphibians.

**Figure 1.**
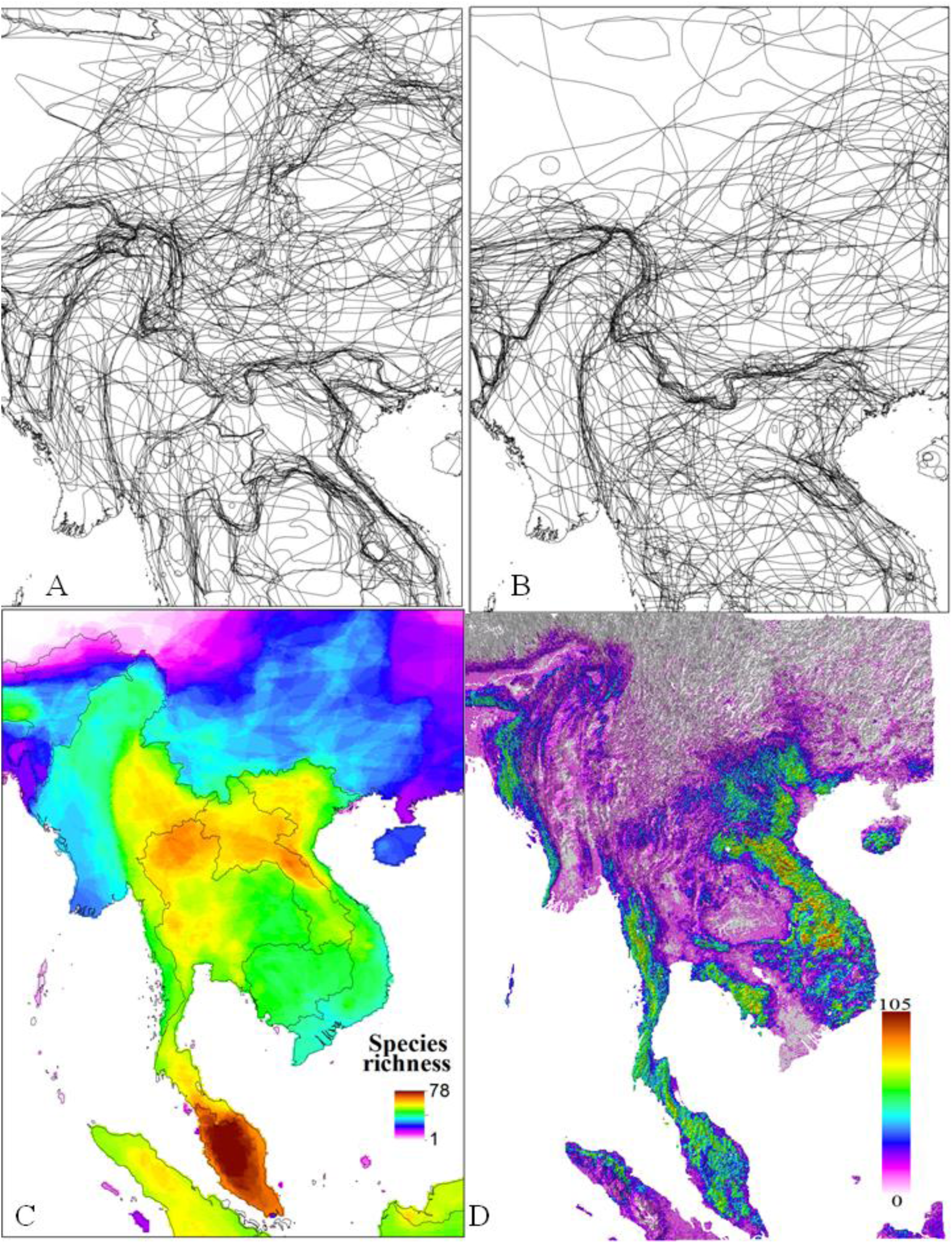
Maps of species distributions and diversity in Southeast-Asia. A-B: IUCN maps of species ranges for A). Rodentia and B). Chiroptera. C shows the IUCN map of bat diversity, and D) the map of bat diversity using predictive modeling.

**Figure 2.**
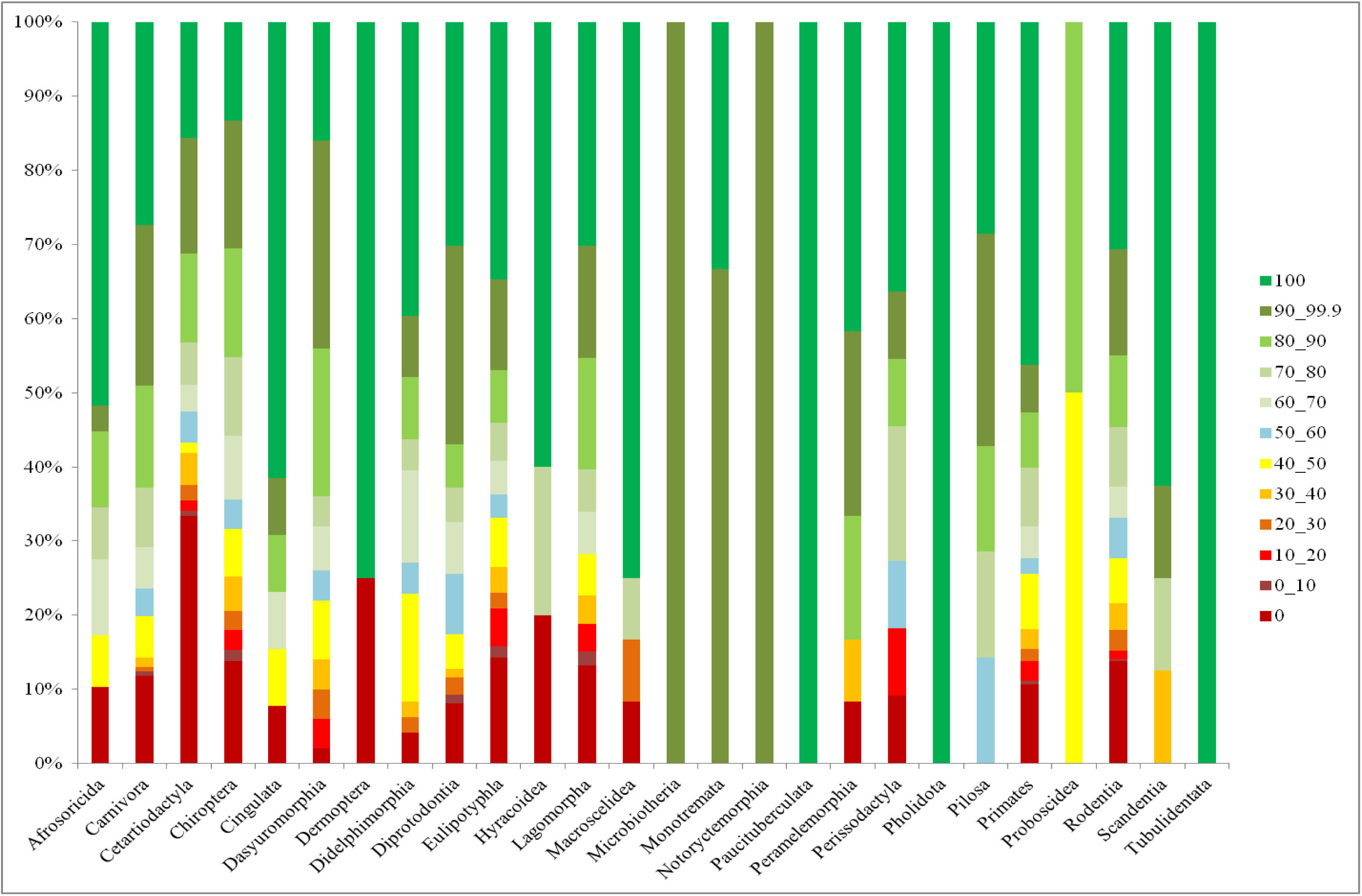
Graph to show the percentage of species each mammalian order with different percentages of known localities falling within the IUCN range map for each species. The size of each coloured portion of the bar denotes what percentage of that order showed the degree of match denoted by the colour of the portion (from 0-0% of recorded localities for each species fell within the IUCN mapped range to 100: 100% of localities fell within mapped range).

Overall Southeast-Asia is the least well biotically mapped region (Figure 3, 60% accuracy on average) followed by Australasia (64.15%), Africa (66.01%), Eurasia (69.07%), South America (70.04%), North America (78.39%) and Europe (81.03%). The Certartiodactyla have the lowest degree of congruence between distribution maps and recorded localities in all regions except North America. Most widely distributed orders showed the lowest degree of accuracy in Southeast-Asia (Certiodactyla-26.42%, Chiroptera-55.67%, Eulipotyphla-55.53%) whereas Rodentia though poorly mapped in Asia (60.51%%) are even less well mapped in Africa (58.33%). In Africa, Rodentia (58.3%), Eulipotyphla (59.3%), and Lagomorpha (56.5%) are all below average accuracy for the region (66%).

**Figure 3.**
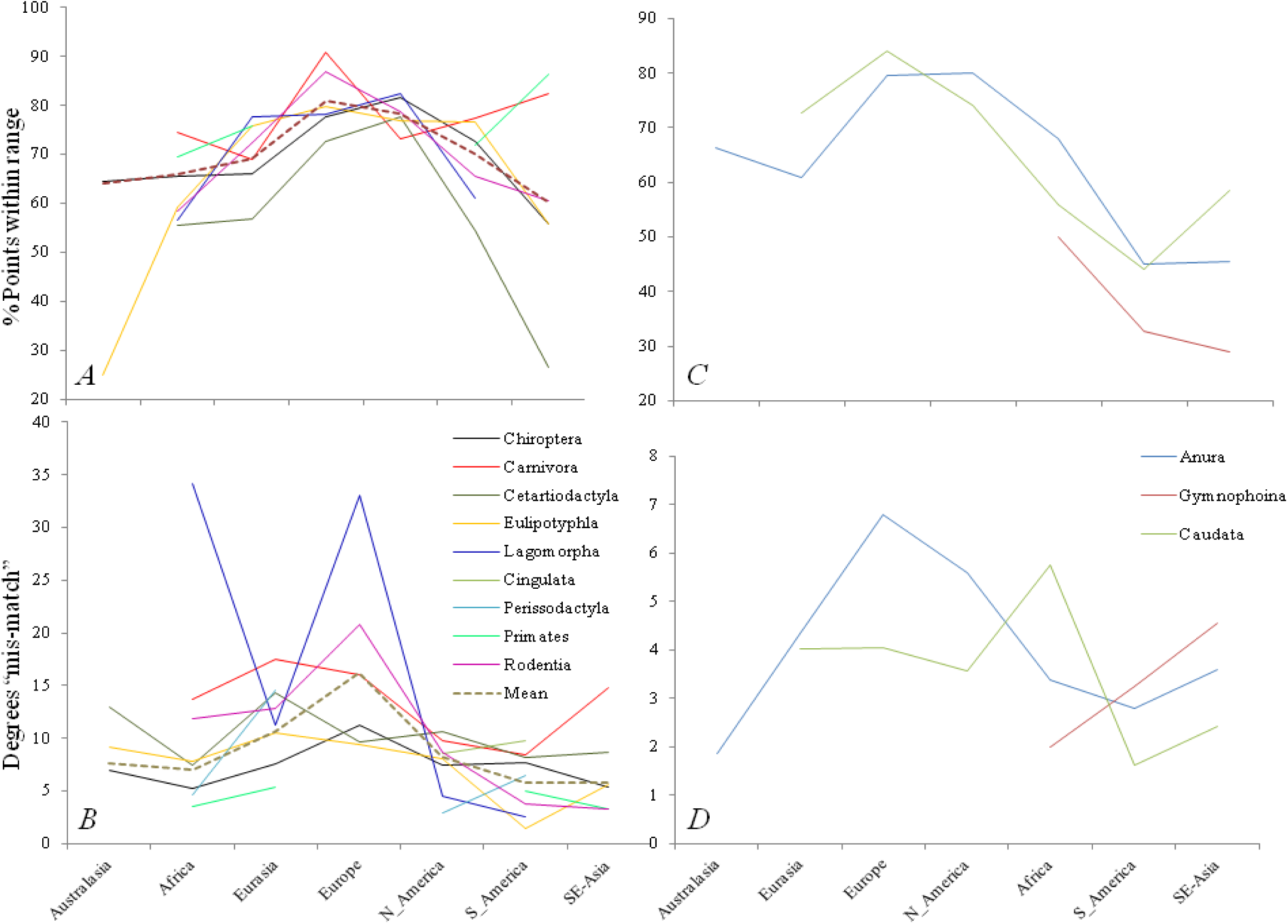
Regional accuracy of IUCN maps for each taxonomic group in different regions. A-B: Mammalian orders, C-D: Amphibia. A and C both show the percentage of recorded points fall within the IUCN range for each species in each zone, averaged by family. B and D show the distance (in decimal degrees) between the recorded and IUCN boundaries of each species range, grouped by family within each region. Greater percentages of accuracy (A, C) mean a higher accuracy, whereas greater distances between recorded and mapped range (in B and D) show a misalignment between species actual and IUCN range.

#### Amphibians

Many of the 2707 amphibian species are represented by a single locality, thus assessments of the number of recorded localities within the range-map for these singletons yields 0 or 100%. Nevertheless, for the 2314 anuran species for which both range and recorded locality data existed 52% of recorded localities fell within the appropriate range, 36% for the 41 Gymnophoina species and 59% for the 352 Caudata species. Removing species for which only a single record exists has little effect to overall accuracy (59, 33 and 66% respectively). South-America and Southeast Asia are the least well mapped region for all three groups (Figure 3C).

### b). Spatial mismatch

#### Mammals

The average difference between recorded and IUCN ranges latitude boundaries showed was 5.03° decimal-degrees (approx 558.33km at the equator), with an average discrepancy of 5.14° at the Southern-most and 4.93° for the Northern-limit. The longitudinal discrepancies were considerably larger at 8.37° overall, 7.78° for the Westerly boundary and 8.96° for the Eastern range limit. Considerable differences exist between taxa (Table 1, and Table S1). Poorly sampled groups. Species poor families (i.e. Tubulidentata, Hyracoidea) show high levels of mismatch, possibly due to undersampling leading to recorded data underestimating species probable range. However for some of these species-poor families (Scandentia, Pholidota) range over-estimates in IUCN maps are also possible, and further research is needed to establish accurate ranges.

Two further forms of error were common in more specious large-ranged species (i.e. 141 Certartiodactyla: 8.79°, 161 Carnivora 12.31°, 11 Perrisodactyla, 7.65°, and 53 Lagomorpha, 8.87°). These groups commonly had IUCN-ranges to the coast, and in Eurasia extensive longitudinal ranges, with little conclusive data to validate these ranges. Many species also showed multiple discrete sub-populations (95 Carnivore and Certartiodactyla species had 313 discrete populations), yet range maps showed contiguous range for the majority of these species.

A further form of error was observed in very specious small-bodied (average: 28.93-214.65g (Jones et al. 2009)) species (Chiroptera 6.9°, Rodentia 6.18°, Eulipotyphla 7.41°, Macroscelidea 6.6°). Many of these species are relatively little studied and little known, thus informed range estimates are difficult and political boundaries have frequently been used as convenient range delimiters (Figure 3-4).

**Figure 4.**
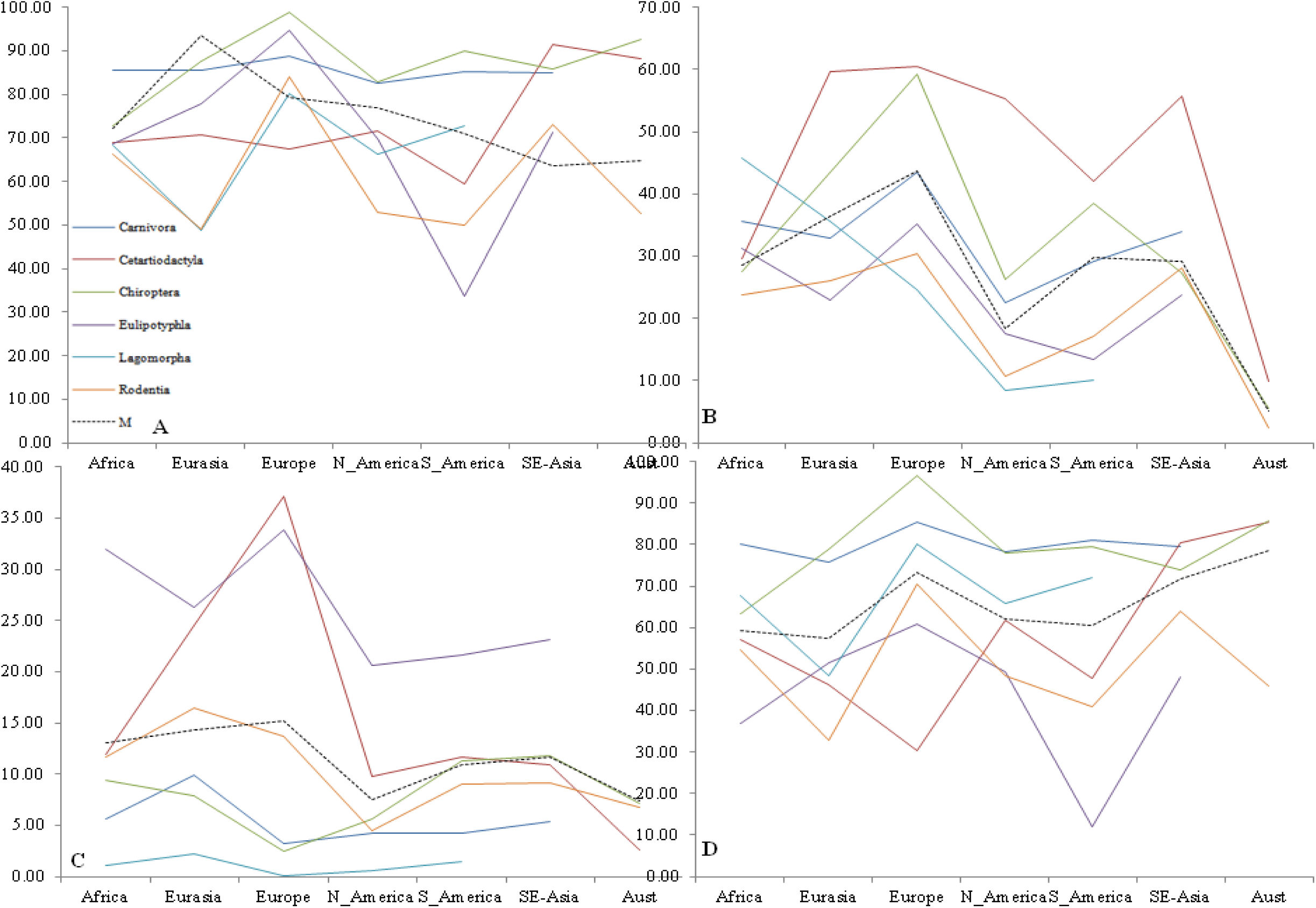
Percentage of IUCN range of each of the six most specious mammal orders (4272 species) which overlaps with political boundaries. A. Percentage of total species range on political-boundaries. B). Percentage of total species range on coastal political-boundaries. C. Percentage of total species range on non-coastal political-boundaries. D. Percentage of non-coastal species range on political-boundaries.

Europe, despite the high proportion of recorded localities falling within the appropriate species ranges has the greatest level of spatial mismatch between recorded and mapped localities (16.23° denoting over-estimates of range size). Contrary to this, regions such as Southeast Asia which have much lower numbers of localities falling within mapped ranges, have a lower degree of spatial mismatch (5.84°), possibly due to more conservative mapping and thus smaller on average range-sizes. Overall the carnivora show the greatest degree of disparity between mapped and recorded distributions, and is consistently above the average degree of mismatch in all zones. Rodentia show a high level of spatial mismatch in temperate and arid regions, but below average levels of mismatch in Southeast Asia and South America (both tropical forest regions). The Lagomorpha are also very poorly mapped relative to collected samples in both Africa and Eurasia.

#### Amphibians

Within the amphibians on a family level the mean distance for Gymnophoina was 0.72°, 3.23° for Caudata and 1.92° for Anura. Within the Anurans the Bombinatoridae had the greatest spatial inaccuracy (at 13.12°) followed by Allophrynidae at 9.84°, whereas all other anuran species had under 2° between recorded and mapped localities. Caudata showed consistently greater distances between recorded and mapped distributions, with 60% of families showing at least 2°, and 30% of families showing over 5° between recorded and mapped distributions (Amphiumidae-5.78°, Proteidae 5.53°, Sirenidae 7.51°). Gymnophoina showed the smallest spatial-mismatch between recorded and actual data, with only two families showing a distance of over 1° (Dermophiidae, 1.42°, Siphonopidae 1.04°). For Anura Europe had the greatest discrepancy (6.8°), whereas in Caudata it was Africa (5.75°) and Southeast-Asia in Gymnophoina (4.57°).

Within the amphibians the Gymnophoina have the lowest degree of accuracy (both in terms of recorded localities within the mapped range, and spatial mismatch) within Southeast Asia (28.89% accuracy and 4.57° mismatch), whereas Africa is much better mapped in both of these regards (50.12%, 1.99°) and South America falls between (32.72%, 3.23°). Within the anurans Europe and North America show the greatest degrees of spatial mismatch (6.8 °, 5.6 °), but as for the mammals they also show most recorded distributions within mapped ranges (79.59, and 79.94%) denoting probable over-estimation of ranges for many species. Southeast Asia and South America showed the smallest number of recorded localities within mapped ranges for anurans (45.54 and 45.07%) but lower spatial mismatch (3.59 and 2.78 °). For Caudata South America shows the lowest number of recorded localities within mapped ranges (44.15%) but also shows the lowest spatial mismatch at 1.61 °, and once again Europe shows the highest classification accuracy (84%), but also a high degree of spatial mismatch (4.044 °, though Africa is higher at 5.75° and has a lower accuracy at only 55.99%).

### c). Species boundaries and political boundaries

To assess the usage of political-boundaries and coastlines as range delimiters (despite changing vegetation in most coastal regions) the congruence of IUCN-range-boundaries and political borders were also assessed for 4272 species in the six most specious mammal orders (Figure 4). Overall political boundaries concurred from 66.44% in the Rodentia (2119 species), to 85.40% in Chiroptera (1141 species), though coastlines make up a considerable proportion of this (71.92% Chiroptera to 41.29% in Eulipotyphla (442 species)). Carnivora (246 species) also have a large percentage of range-boundaries along coasts (82.05% of range-boundaries follow political boundaries, 72.69% coastal). For non-coastal boundaries Europe has the highest match between species range-boundaries and political-boundaries (43.63%) and North-America the lowest (18.45%). Certartiodactyla (232 species) and Chiroptera consistently have the highest percentage of concordance between non-coastal range-boundaries and political boundaries (Certartiodactyla: Africa: 29.52% to Europe: 60.58, Chiroptera: 26.35% in North-America-59.14 in Europe) followed by Carnivora (North-America 22.56-43.47% Europe). Further regional breakdown is available in the Appendix S2, and Table S1.

## Discussion

The level of disparity between recorded and mapped ranges is startling ranging from hundreds to thousands of kilometers, as high levels of error and inconsistency between recorded and IUCN mapped distributions are common across all taxa. Within the mammals four main types of error in mapped ranges were common for different taxonomic groups. Range-overestimation (observed records exist within mapped range, but there is a considerable distance between recorded and IUCN outer boundaries) is common for all taxa. This form of error is common across taxa, but particularly for rare species, with limited number of samples. One of the major implications of this form of error is to assume a species occupies a much larger range than it occurs over, as a result the species may be assigned a less “at risk” IUCN redlist status than actually warranted. As a result of having a lower redlist status the species is less likely to receive funding from grants, or to have it’s actual range protected, thus on a species level this form of error actually impairs conservation efforts.

The second form of error, which was observed most in Carnivora, Certartiodactyla and other taxa with larger bodies (over 1kg on average) and ranges, is a longitudinal overestimation of range, in some cases by thousands of kilometres where no databased records for the species exist. A number of these species are mapped as having extensive ranges (often circumpolar), such as *Gulo gulo*, yet mapped distributions show restricted regional populations (in China, Europe and North-America (Zhang et al. 2007), with limited published data and anecdotal data on distributions outside these areas. Genetic approaches when used are likely to reveal that these species only occur over a fraction of the range that they are mapped to occur over, and that the populations which do exist have limited geneflow and thus require separate conservation efforts to adequately conserve. This form of error relates to the third type of error, which is also common in the same large bodied taxa. In this form of error large contiguous ranges are mapped to exist when infact a number of smaller populations are mapped, and may have existed as separate populations for extended periods of time (i.e. since the last glacial maximum). Many of these large bodied taxa showed upto six separate populations (i.e 1-4 in *Canis aureus, Felis chaus, Lontra longicaudis)*, which though also possibly showing the patchiness of databased research in some instances are likely to at least for a large proportion of species refer to species actual distributions. As many of these populations are separated by clearly unsuitable areas they are likely to be genetically distinct to varying degrees, and therefore mapped ranges that show an area as contiguously suitable rather than patchy not only overestimates species range, but underestimates the vulnerability of distinct populations. The lack of recognition of these populations also means that whereas connecting areas (i.e. through afforestation) is a sensible approach in many circumstances, it makes little sense to connect possibly long separated populations, which may be adapted to local conditions, or to relocate individuals between very distant and unconnected regions. The IUCN guidelines mentions recording these details, yet they rarely appear on maps, but can change the way that conservation for these species is planned.

For smaller mammals (less than 250g on average (Jones et al. 2009)) a fourth, and type of error is common, especially where scant ecological data is available for any given species. In this case political, or other convenient boundaries may be used to delimit species ranges. Political-boundaries are clearly visible on IUCN-maps of Southeast-Asia, where the popular perception that China is not part of Southeast-Asia has led to a considerable number of small-mammal species (possibly hundreds, Figure 1) having the Chinese political border as the Northern-limit of their mapped range. Political-boundaries should not be visible on biodiversity maps, except where differential management practices in neighbouring countries (i.e. Thailand:Lao-PDR) has led to different levels of forest coverage or hunting pressure in areas which may once have harboured very similar species. This form of error, in addition to meaning that areas of biodiversity are likely to be entirely miss-classified (as these groups include the most specious and under-studied groups, and mapped ranges of species are likely to be developed on the basis of relatively little empirical data). In Southeast-Asia the mammal diversity maps from the IUCN (i.e. Chiroptera: Figure 1 & 3) do not match current forest cover, or adequately reflect current research on Southeast-Asian patterns of diversity. Thus any decisions based on these maps will result in conservation of areas of dubious diversity at the expense of areas with greater diversity, as evidenced by the striking contrast between the diversity maps shown by the IUCN (Figure 1C) and through species distribution modelling (Figure 1D). Areas such as Southern-China, and other regions which the IUCN classes as low diversity will find it difficult to find funding, easy to obtain permission to develop and are unlikely to be protected, which may have devastating implications for biodiversity. Thus rather than assisting conservation efforts, the IUCN maps may actually hinder effective conservation, through inaccurate status assessments of species vulnerability (as outlined above), and incorrectly mapping out biodiversity patterns (which forms the basis of protected area planning on a regional and global scale (1-5).

Though the IUCN guidelines should prevent the aforementioned errors, it is clear that for many species their distributions are not accurately reflected at present. As a result of these errors (21.4% of mammal records distributions fell outside mapped ranges and 47% of amphibian distributions) scientists and managers need to re-evaluate how we map out species distributions, and provide more informed and relevant information for the protection of these species and regions. This can be more reliably done, without the need for huge amounts more data, as within the last two decades the volume of data available of all forms has increased tremendously.

The IUCN sees the maps as an essential component of redlisting-assessment, for visualising and understanding ranges and as a basis for conservation decisions. However though the online training course advocates a sensible method of starting with distribution data (https://www.conservationtraining.org), and using this as a basis for the range map, there is still significant human intuition required, which is reflected by the usage of political boundaries and level of errors found. The training for development of species maps for terrestrial, marine and freshwater systems is expected to take 30 minutes in total and at present the redlist syllabus lacks explicit GIS training. Thus though the IUCN values these maps, training and testing need to be dramatically altered in the future, to prevent the current level of error being continued.

### Solutions

One simple alternative approach, which may refine the IUCN range-maps and improve their spatial accuracy is to refine current IUCN range-maps with actual vegetation-cover data. If the species of interest is known to be associated with forests for example, then non-forested areas could very rapidly be removed from the IUCN range-map of the species. In a region such as Southeast-Asia, even this simple step would transform the current biodiversity map (as especially for small mammals the range-maps may be created with little regard for land-cover or topography). This is a minimum step which should be considered before any further usage of IUCN distribution maps.

A better method is however now available, which relies on the ever increasing volume of available data on species and their distributions. This study utilised cleaned GBIF data because it is available to researchers globally and allows accurate modelling of biodiversity. By pairing species distributions with the various eco-geographic conditions which prevail it is possible to continuously map a species distributions *(6)*, and unpick the factors underlying the distribution of any given species. Many species which exist within museum collections are not yet included in IUCN mapped species, thus the usage of existing data on species distributions actually increases the number of species which can be mapped and used in planning analysis. Even for species which have only been collected once, it is possible to extract environmental data using whatever limited data exists, and map out areas which meet those conditions for further research or conservation processes. Given that the IUCN runs workshops to train assessors of species vulnerability, it would be a fairly simple step to teach them how to develop and test species distribution models, which make use of the abundant data now available on species distributions and the environmental factors which may drive the species distribution. Maps created on these grounds would much better reflect species ecology, in addition to lending insights into habitat requirements which may provide further insights which facilitate effective conservation.

However, a poor model of a species distribution is no better than a poorly informed map of a species occurrence, and researchers and managers often wish to know the whereabouts of a species without the time or knowledge to produce a species specific model of occurrence for that species, and it is probably most often these individuals who make use of IUCN maps in decision making and planning. Yet there is still a viable alternative to IUCN maps, which makes use of available data to produce a map of species occurrence without the need for lengthy analysis and data preparation.

The Lifemapper website (lifemapper.org) utilises GBIF data in combination with environmental data to enable any individual to map a species distribution, and download the resulting map in various formats. The outcomes of such analysis can be used as a viable alternative to the current IUCN maps, as they can be produced and accessed by any individual and are based on empirical data rather than human impressions of where a species might be. Though these measures can also be improved, if users could evaluate the accuracy of spatial data in order to remove any suspect or erroneous records (as the atlas of living Australia does: spatial.ala.org.au, code.google.com/p/ala-dataquality). Given the flexibility to map any species, with data that can be visualised, interacted with and checked would provide the reliability needed to map out and best protect the biodiversity of this planet into the future.

In an age of peer-review, big data and sophisticated approaches to model various ecological phenomena, we need to use more empirical approaches to respond to ecological issues. This is nevermore the case than when such data is used to allocate funding, prioritise species and areas and in attempting to understand global distributions of species across the planet, and when we now have the data and the technology to make better decisions for global biodiversity. The current usage of IUCN maps in global biodiversity and conservation assessments will lead to poor decisions, especially in poorly mapped regions such as Southeast-Asia, thus the use of better methods to map biodiversity are essential before any further evaluations of biodiversity patterns and protection. The IUCN aims to facilitate conservation globally, but to do so requires a more empirical approach, which more accurately reflects current diversity and helps inform and direct conservation across the planet.

## Appendix S1

### Supplementary methods

*a): Data cleaning.* Distribution data was cleaned by sorting species according to latitude and minusing each latitude for a given species from the closest record for that species, these values were then highlighted using conditional formatting, as were the actual latitudes of occurrence. Chronologically neighbouring species were removed from this sorting using the “match” function in excel. Any point falling a significant distance from neighbours (more than 5°) was given a numerator following the species name, or excluded altogether (if the record clearly fell on a different continent different biome from the bulk of data for the species). Data was then sorted by longitude for each species and the process repeated. Ambiguous records for any genera were assigned to species when their latitudes and longitude values were within 0.01° of a record for a species of the same genera. Duplicate records for a species at any particular locality were removed, to remove biases caused by intensive surveying at any particular locality relative to more sparse, or opportunistically surveyed regions. Species which clearly formed a number of separate populations were also enumerated, as especially in such species (including large numbers of Certartiodactyla and Carnivora) the continuous IUCN range-maps greatly overestimate the species actual range, and many of these populations may actually be distinct subspecies and are clearly unable to interbreed. The distribution of all sub-populations were checked within ArcGis and possible introduced, captive and invasive samples removed from further analysis.

Once this was complete data was uploaded into ArcView 10.1, and convex hulls formed for each species using the Minimum bounding geometry function to assess the presence of any outliers (i.e single records falling on different continents) for any of the species present for each taxonomic group.

*b).* Once exported to excel any points with no intersecting IUCN polygons were removed. The match function was then used to find matching species for point data with intersected polygon data, unmatched point data removed and the “countif” function used to identify the total number of points for each species fell within the appropriate range polygon. This was repeated with any sub-populations for species (noted by species name followed by a number). The total number of distribution points for each species (from cleaned data) was uploaded in ArcView with an arbitrary latitude and longitude and the species field joined to the species “point in polygon data”. The resulting table for each taxa was then exported and the percentage of distribution points falling within the appropriate range polygon calculated for each species.

*c)*. To examine the spatial mismatch between actual distribution records and IUCN range-map we extracted the maximum and minimum latitudes and longitudes of occurrence from the distribution point data using the “summarise statistics” tool in ArcView. To determine maximum and minimum range dimensions from IUCN range-maps the “convert vertices to points” tool was used, followed by summery statistics. Summary statistics were then used to find the dimensions of each species range, and these dimensions compared to those from actual distribution data.

*d)*. The concordance of species range boundaries and political boundaries were examined for the six mammal orders which spanned at least five of the biogeographic zones, giving a total of 4272 mammal species (2119 Rodentia species, 1141 Chiroptera species, 442 Eulipotyphla, 246 Carnivora, 232 Cetartiodactyla and 92 Lagomorphia). To examine the concurrence between political boundaries and species ranges the IUCN range-maps for each species were converted to points using the “vertices to points” tool in ArcGis (producing upto 7803124 points for each of the six orders). Using the country boundary data (http://www.diva-gis.org/Data) the spatial join function of ArcGis was used to intersect each species range-map with any nearby political boundaries. A 25km buffer each side of the political boundary was specified, to capture the ranges of any species which “approximately shadowed” a political boundary. The process was repeated using the output of the political boundary intersection with a coastal boundary (http://www.ngdc.noaa.gov/mgg/shorelines/gshhs.html). The total number of points which formed each species range boundary was compared to the number along the political boundary, the coastal zone and related measures to determine the degree to which the political boundary was used as the range delimiter of any given species. The results were then broken down by order and by region, to look for spatial biases in the usage of boundaries to define range limits. Zones with relatively few political boundaries (i.e. Australasia) are unlikely to utilise these boundaries heavily, whereas areas such as Europe which have a relatively large coastal zone will reflect the usage of coastal to non-coastal political boundaries through the proportionate discrepancy between total non-coastal political boundary and percentage of non-coastal range-boundary which overlaps with a political border.

## Appendix S2: Commentary

### A-Range coverage by taxa

For mammals at the family level many families only had a single or small number of species, and of the fourteen families where an average of below fifty percent of recorded localities fell within species range-maps, six were Rodentia, three were bats, and two were cetartiodactyla (in addition to one carnivore, one Peramelemorphia and one Dasyuromorphid family). On a family level at least ten families (with at least three species in each) have at least 50% of their species showing under 50% of known localities falling within the mapped range for those species (*Anomaluridae, Chrysochloridae, Ctenomyidae, Echimyidae, Erethizontidae, Galagidae, Moschidae, Rhinopomatidae, Thyropteridae, Tragulidae*-See supplementary information figure one). Of families with at least five species only one (Dinomyidae: Rodentia, five species) was only recorded entirely outside its mapped range, Ctenomyidae (also a rodent family, 13 species considered) was the next least well mapped with only 37.8% of recorded localities falling within the mapped range. One further family of over five species also had under 50% of records within the IUCN range, the cervidae (cetartiodactyla, 29 species) had only 47.69% of localities falling within their mapped ranges. A further twenty of the 128 mammal families considered (11.72%) had 50-60% of recorded localities falling within the appropriate mapped range, 15.63% of families had 60-70% of localities within the mapped ranges, 22.66 % had 70-80%, 18.75% of families had 80-90% of localities within mapped ranges and the remaining 20% of families had over 90% of records within the mapped range.

If Amphibian orders are split into their constituent families (49 for Anura, 9 for Caudata and 8 for Gymnophiona) there is a marked difference in the sampling intensity and accuracy for different taxa. When singletons are removed from the analysis accuracy in the Gymnophiona varied from 0 in the Scolecomorphidae, and the Herpelidae to 72.99% in the Dermophiidae, and the average accuracy for families was 34.04%. The Caudata are considerably better varying from 62.43% (Plethodontidae) to 100% in the Cryptobranchidae, with 44% of families showing an accuracy from 60-70%, 33% of families between from 70 to 80% accuracy and all further families over 90% accurate. For anuran families the average was 62.79% of records falling into appropriate mapped ranges, however this was very variable (standard deviation: 24.76). Other than the one anuran family with over one sample but showing no locations falling within mapped ranges (Nasikabatrachidae) the next poorest family was Alsodidae, with only 13.33% of occurrences falling within mapped ranges. For the anurans (with singletons removed) 26.67% of families had below half recorded occurrences falling within mapped ranges, 42.22% between 50 and 75% accuracy, 13% from 75-90% accuracy and 17.78% of families had over 90% of distribution points falling within mapped ranges.

### B-Spatial mismatch by taxa

Poorly studied groups will naturally show a greater discrepancy between recorded and mapped ranges due to undersampling, and thus small poorly studied groups such as the Tubulidentata may be due to lack of recorded specimens rather than poor mapping. The same is likely to be true for the next most spatially mismatched mammal group, the Hyracoidea (average 26.01° discrepancy, but 74.3% of localities fell within mapped range) which has a single well mapped species (*Dendrohyrax validus*: 2.75° discrepancy) and four poorly mapped species across a large poorly studied region. Scandentia was overall the third most mismatched taxa (18.75° discrepancy) and with the low proportion of recorded localities falling within the mapped range this is liable to represent range underestimates and distribution errors for most species. Pholidota (13.66° discrepancy), like Pilosa and Dermoptera (8.12° and 5.59° discrepancies) in addition to many of the smaller marsupial groups also represent undersampled rather than poorly mapped species, as all have the majority of locations falling within mapped ranges despite large discrepancies between mapped ranges and recorded localities. However for all these potentially undersampled groups there may also be a large degree of range-overestimation, where the IUCN mapped range is much greater than the actual distribution of each species, therefore causing a large discrepancy between recorded and mapped range, whilst still showing that at least part of the mapped range is correct.

The next most poorly mapped orders have considerably more species, the Carnivora (161 species, 12.31°), Lagomorpha (53 species, 9.87°) and the Certartiodactyla (141 species, 8.79°) and the Perissodactyla (11 species, 7.65°). Thus though the discrepancies between recorded and mapped boundaries may represent poor sampling for some species, it is liable to represent genuine IUCN map errors for the majority of cases (concordant with the percentage of localities outside recorded ranges for these taxa, 24.9, 47.4, 29.4 and 26.6% respectively). These groups are generally large and wide ranging species, whilst not typically difficult to classify or recognise, thus errors in maps are likely to result from the over-simplification of range shape (these groups had the greatest number of sub-populations (313 populations of 95 species of carnivore and Certartiodactyla)), and poorly informed range limits based on assumptions of distributions in undersampled regions.

The next rank of taxa in terms of mismatch are generally smaller, more difficult to accurately classify and frequently poorly studied, these order have a spatial mismatch from 6.18° (Rodentia) to 7.41° (Eulipotyphla) (in addition to Chiroptera at 6.9°, and Macroscelidea at 6.6°). For these taxa political boundaries are most likely to be used as convenient range delimiters for significant portions of species (see Figure 1).

The final group of orders with a spatial mismatch range of 0.85° (Microbiotheria) to 5.44° (Didelphimorphia) and is largely composed of marsupials, in addition to the Monotremata and a select number of other orders. Many of these groups occur only within Australia, and are tightly linked to bio-climatic zones, thus limiting the margin of error possible, which is supported by the high concordance between recorded localities and mapped data (75.55-99.3%). Unsurprisingly the order with the higher degrees of inaccuracy are the Didelphimorphia which fall outside Australia, though the two orders within South-America (Paucituberculata and Microbiotheria) are both fairly depaupaurate and well known in addition to occurring in a fairly limited region, and therefore have relatively low levels of inaccuracy (2.65° and 0.85° respectively). The four remaining marsupial orders considered (Diprotodontia 2.82°, Notoryctemorphia 2.84°, Dasyuromorphia 3.5°, Peramelemorphia 4.98°) and the Monotremata (2.36°) are all confined to Australasia and thus have relatively low inaccuracy.

This final group also contains three therian groups, the Proboscidea (3.08°), the primates (3.91°) and the Afrosoricida (3.36°). The primates are generally a fairly well studied group relative to their diversity, as are the Proboscidea, which could explain the relatively low degree of mismatch between observed and mapped range boundaries. The Afrosorcida is largely confined to Madagascar, limiting the average spatial mismatch for a significant number of the group.

---

**Table S1**

**Table 2.**
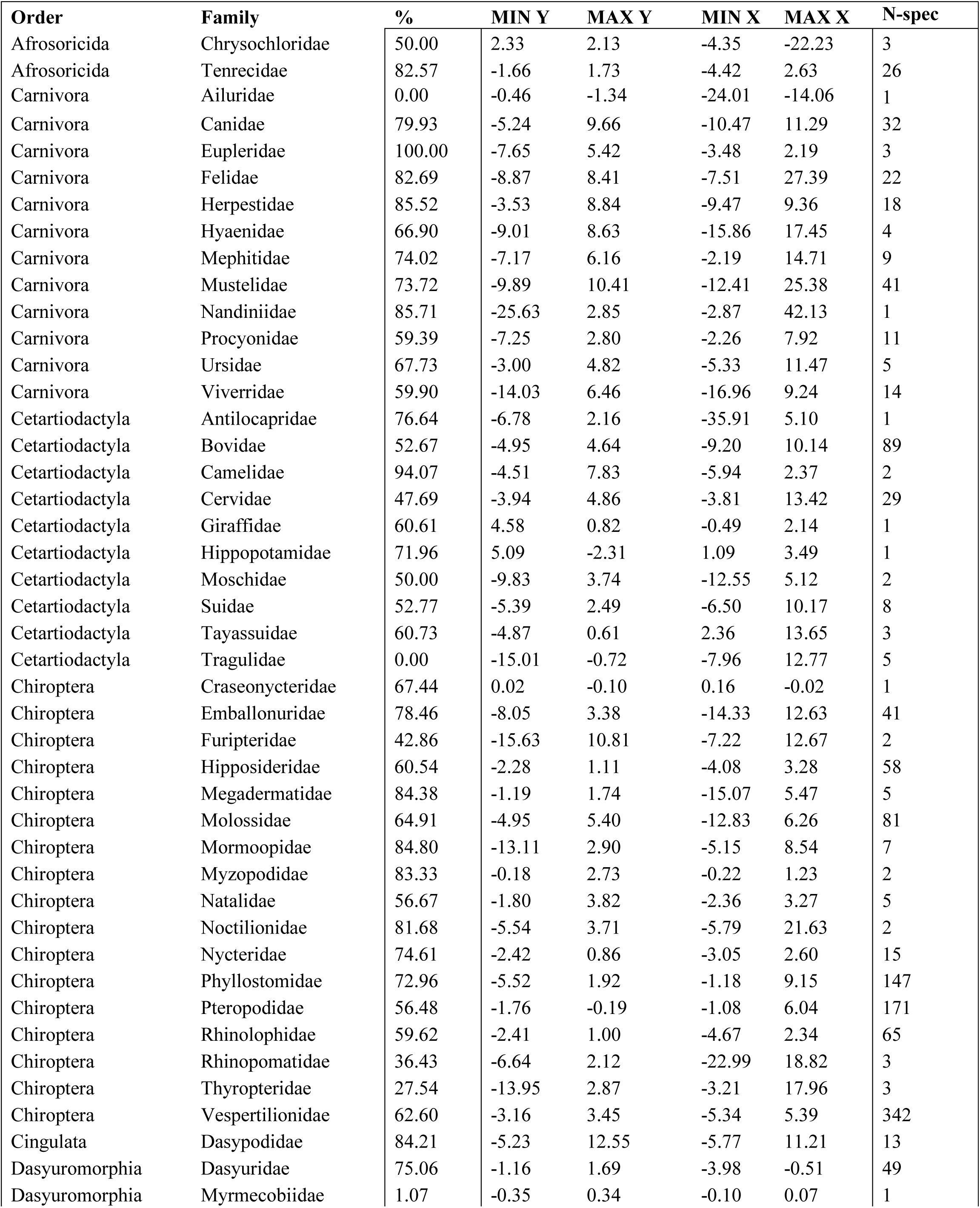
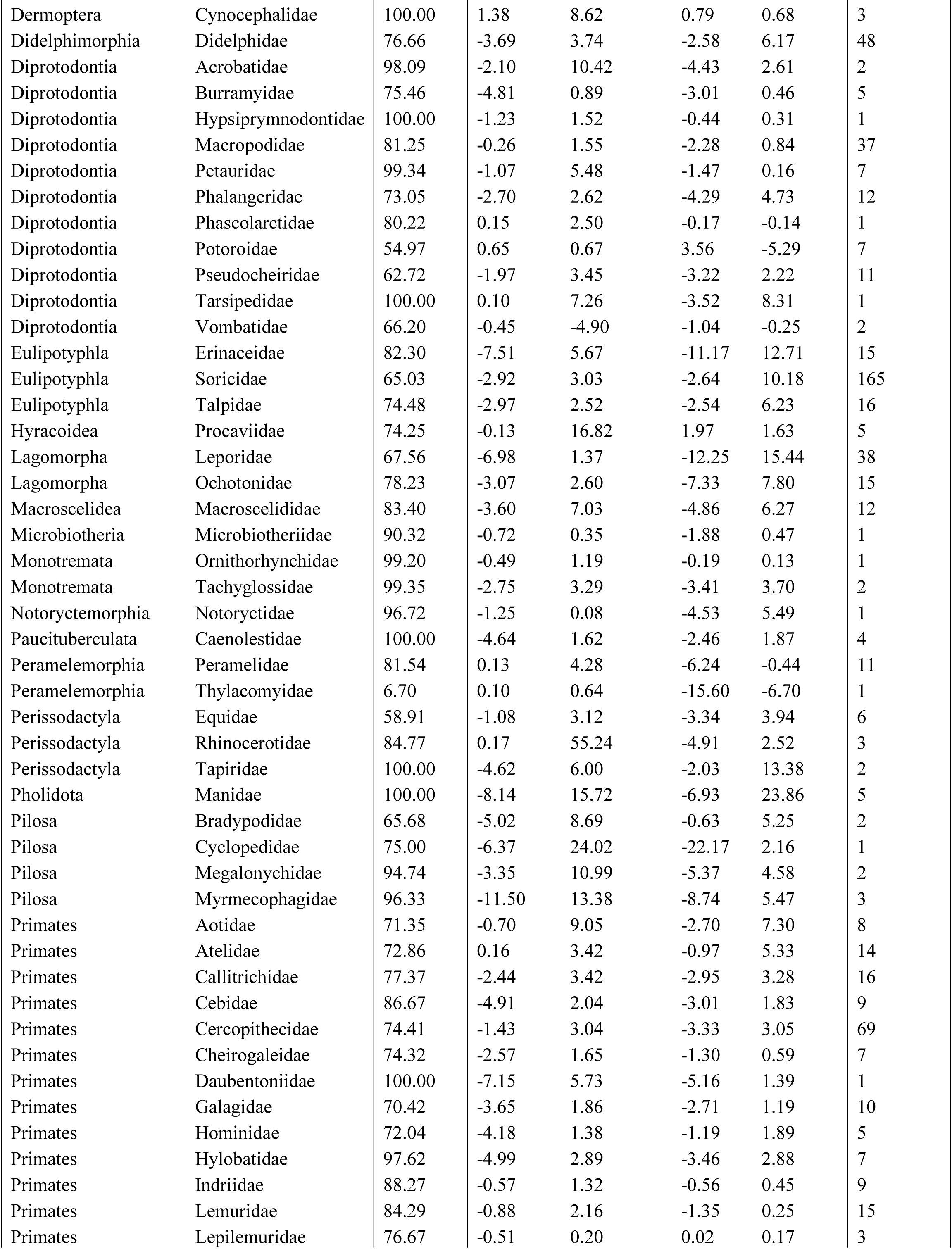
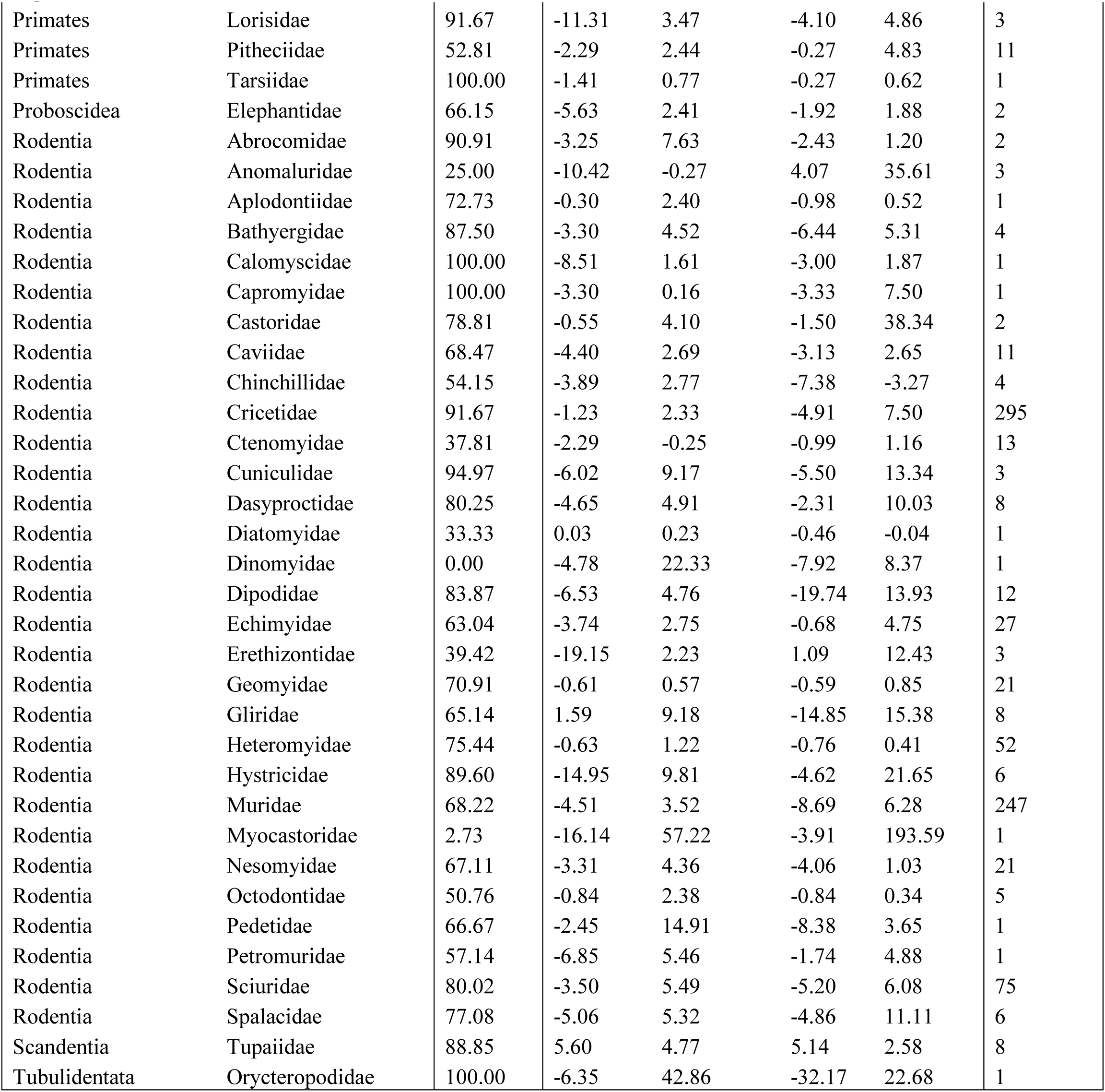
Accuracy assessment for each family, both for degree of congruence between mapped ranges and recorded range boundaries for each species, and for the percentage of recorded localities for a species fall within the appropriate range map.

